# MOTUM: a system for Motion Online Tracking Under MRI

**DOI:** 10.1101/2025.07.26.665107

**Authors:** Federica Bencivenga, Michelangelo Tani, Krishnendu Vyas, Federico Giove, Steve Gazzitano, Gaspare Galati

## Abstract

Attempts to implement realistic body-environment interactions during functional magnetic resonance imaging (fMRI) experiments have developed expensive, hardly reproducible, and task-specific setups. Here, we introduce MOTUM (Motion Online Tracking Under MRI), a novel system that combines real-time kinematic tracking with immersive virtual reality, enabling participants to perform naturalistic movements inside the scanner. As a proof-of-concept, we tested MOTUM during a reach-to-grasp task with and without visual feedback of one’s hand (N = 7). The system achieved high-fidelity motion tracking, induced an intense immersive experience, evoked expected sensorimotor brain activations, and maintained high fMRI data quality. Standard fMRI control metrics were below the critical threshold in 99% of volumes, indicating that participants’ arm movements had minimal impact on head motion and data quality. While hand movements had little to no effect on brain activity, arm movements resulted in sparse spurious correlations that were easily controlled for. Critically, MOTUM allowed us to extract rich kinematic indices and link them directly to brain activity on a trial-by-trial basis. Parametric modulation analyses revealed that natural variations in movement dynamics significantly influenced neural responses in parietal, frontal, and occipital regions. In sum, MOTUM is a robust method to study motor control and beyond, enabling a new class of fMRI experiments that bridge ecological realism and experimental control, pushing current neuroimaging research towards real-life neuroscience.

## 1. Introduction

Biological motion offers some of the richest and most reliable measures of behavior. Nearly a century ago, Bernstein (1927) proposed that kinematic tracking could provide a unique window into core brain mechanisms. This insight aligns remarkably well with contemporary perspectives, as we now think that brain circuitry primarily evolved to generate adaptive, complex movements (Krakauer et al., 2017), with other sophisticated cognitive functions remaining fundamentally grounded in sensorimotor processes (Pezzulo & Cisek, 2016). Under this perspective, research on sensorimotor integration constitutes a preferential route for understanding the complexity of the brain (Cisek, 2007).

Still, body-environment motor interactions can be acknowledged among the most theoretically and methodologically challenging topics in neuroimaging research. The richness of the motor behaviors driving our exchanges with the environment is complex to replicate in experimental setups, whereby the necessity of controlling confounding variables leads to studying motor control under simplified, often impoverished conditions, with movement being paradoxically constrained. A growing field trend advocates for studying more naturalistic behaviors under controlled experiments, achieving a trade-off between ecological behaviors without systematic control and constrained laboratory setups (Cisek & Green, 2024). The recent advances in virtual reality technologies, such as wireless VR headsets, trackers, and controllers, allow us to meet this criterion and, over the past decade, have significantly boosted research on naturalistic behaviors (Maselli et al., 2023).

An even more formidable challenge is to conjugate the study of naturalistic behaviors with neuroimaging techniques like functional magnetic resonance imaging (fMRI) and magnetoencephalography (MEG) to investigate the neural correlates of cognitive and motor functions in humans with high spatial resolution. Combining MEG with optically pumped magnetometers (OPMs) has significantly boosted the study of naturalistic behaviors (Xia et al., 2006; Boto et al., 2016; Brookes et al., 2022). Beyond being motion-robust, OPM-MEG also provides data with higher quality than traditional MEG, with signals that are larger in amplitude and better spatially localized (Brookes et al., 2022). This is not the case of portable, ultra-low field MRI scanners (Sarracanie et al., 2015), which are beneficial in emergency or ambulatory care but not suitable for research purposes, as they do not achieve the same quality of data as high-field, non-portable MRIs. As such, studies on naturalistic body-environment interactions using fMRI are still lacking.

Although these limitations affect research on many topics (body ownership, social interactions, spatial navigation), here we will focus on the domain of visuomotor functions, namely the mechanisms through which objects’ properties are extracted by visual inputs and turned into motor actions (e.g., pointing, reaching, or grasping). This is an excellent example of how the grounding theories on motor control and object interactions have relied on simplified experiments detached from real-life situations. Indeed, most of our knowledge of such behaviors comes from experiments where participants reach simplified targets, e.g., dots in the space, or grasp ad-hoc created objects with simple geometrical configurations (e.g., manipulanda). Most studies have adopted workarounds such as projecting 2D visual stimuli on a mirror in the scanner, instead of presenting real 3D objects. In these studies, participants were required to execute pantomime movements pretending that the images showed real objects near their hand (Przybylski & Króliczak, 2017; Sulpizio et al., 2020). However, other studies have underlined that action representations, especially in high-order cognitive (e.g., posterior parietal) areas, are affected by the realness of the objects to interact with (Króliczak et al., 2007).

Others have attempted to use real objects within MRI, which is not exempt from issues. Culham’s group has used a “grasparatus” (Culham et al., 2003), implemented by tilting the torso of participants by 5 degrees using a shallow ramp and a 20-30° tilt to the participant’s head using foam. With this apparatus, subjects could see their hand movements and interact with real objects located on a bracket above the MRI table. However, additional devices must be set up to widen the range of objects to be presented. For instance, Brandi and colleagues (2014) have used a “tool carousel”, i.e., a box with six compartments that can hold different objects and be turned around its central axis; Nowik and colleagues (2019) developed a new version of the “grasparatus”, consisting of two rotating drums connected with side panels and a conveyor belt, on top of which the objects were mounted; Knights and colleagues (2022) have used a turntable. In many cases, objects have been created using a 3D printer. A common feature of all these experiments was the presentation of a simplified scenario, lacking the complexity of everyday life. While virtual reality can account for that, to our knowledge, no attempts have been made to integrate VR and motor functions during fMRI exams.

In this proof-of-concept paper, we introduce a methodology to overcome current limits in functional neuroimaging of motor control by using virtual reality combined with an MRI-compatible motion tracking system. We label this method “MOTUM” (Motion Online Tracking Under MRI). Virtual reality makes it possible to create naturalistic environments with a high degree of control, fitting the idea of realistic but still manipulable scenarios. Using MOTUM, participants can act in these environments using a virtual avatar, whose movements are created and updated according to the real-life position and rotation of the participants’ body parts and displayed in a first-person perspective. This is achieved using MRI-compatible motion tracking devices broadcasting information into a VR model.

To test MOTUM, we developed a reach-to-grasp visually guided task whereby we isolated the effect of visual feedback on one’s movement. Participants (N = 7) were asked to grasp narrow or wide parts of a custom 3D virtual object through a precision or power grip. In separate conditions, participants grasped with online feedback of the hand movement provided by MOTUM or without visual feedback. Importantly, we collected and analyzed kinematic data under the two conditions and incorporated them into the imaging analysis pipeline, demonstrating a set of interesting, though preliminary, correlations between kinematic parameters of the reaching movements and regional brain activity in sensorimotor cortical regions.

While describing the results of this pilot experiment as a proof-of-concept of how MOTUM can overcome the limited landscape of body-environment interactions feasible in the MRI setting, we will also discuss the potential of this system to boost real-life neuroscience in the broader frame of body representation and body-environment interactions.

## 2. Methods

MOTUM consists of three main hardware components, complemented by dedicated software modules (Figure 1): a set of cameras that track the movements of the upper and lower limbs; an MR-compatible glove with sensors monitoring finger flexion and extension; and a pair of MR-compatible binocular goggles that present a stereoscopic (3D) virtual reality scene to the participant.

**Figure 1.**
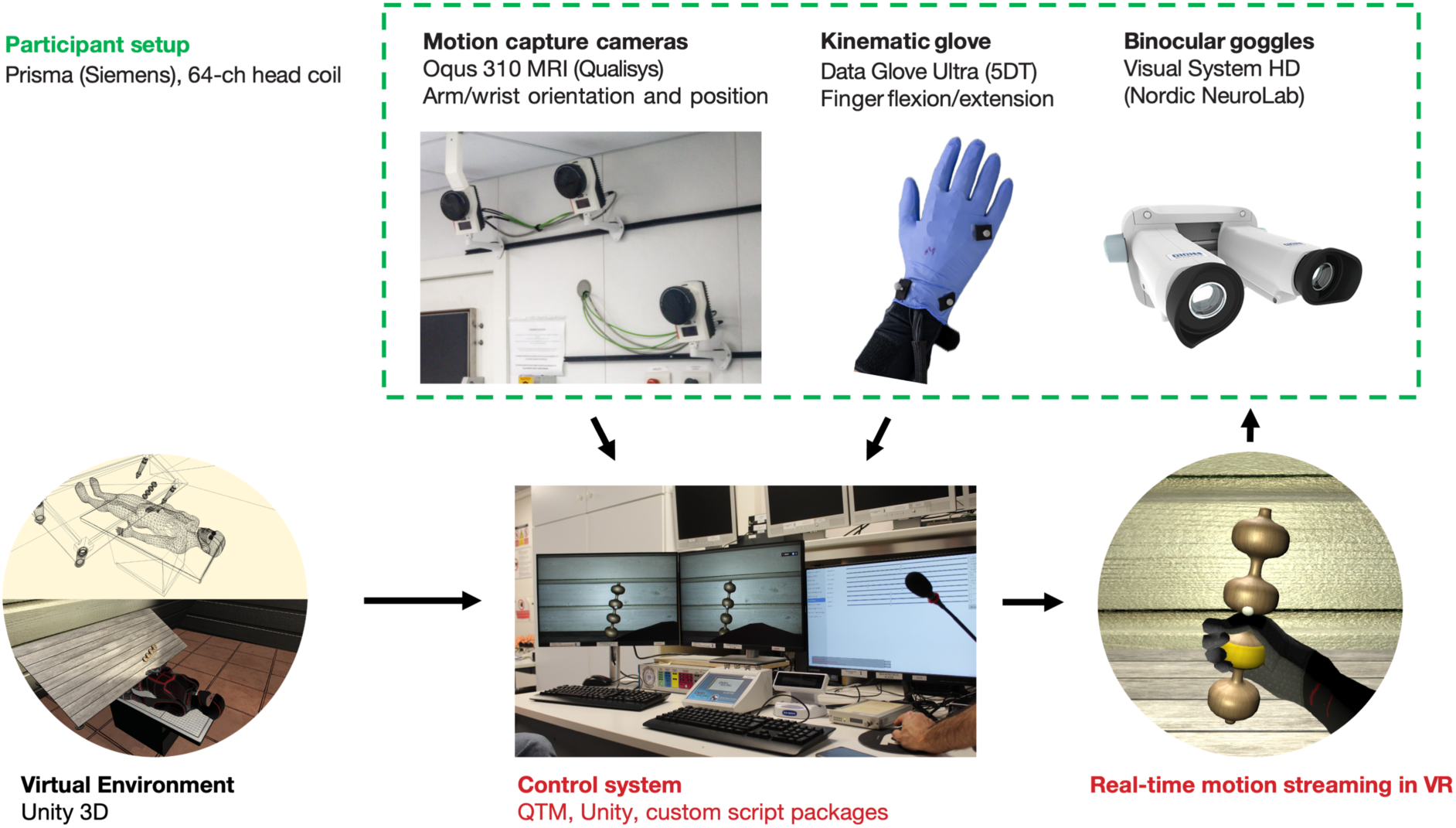
Hardware and software components of the MOTUM system. A participant performs a reaching-to-grasp action with her right hand within the MR scanner. At the same time, a set of MR-compatible motion-capture cameras tracks right forearm movements, and an MR-compatible kinematic glove tracks right-hand finger flexion and extension. A control computer processes real-time kinematic data and feeds it into a virtual environment, animating a humanoid hand in real-time. A first-person perspective is then projected to MR-compatible binocular glasses for an immersive experience.

During the fMRI session, markers placed on the participant’s limbs are tracked to estimate the joint movements of a humanoid skeleton. These data are then combined with finger movement data from the glove and streamed in real time to control a custom-made avatar within a virtual environment. The scene is presented from a first-person perspective through the binocular goggles. Kinematic data are also saved with behavioral data and synchronized with MR data acquisition.

In the present implementation, the camera system tracks markers placed on the participant’s right arm while the participant performs reach-to-grasp actions towards a visual target.

### 2.1 Apparatus

The system described here has been installed at the Neuroimaging Laboratory, Santa Lucia Foundation, Rome, Italy, equipped with a Magnetom Prisma (Siemens, Munich, Germany) 3T scanner.

Three electromagnetically shielded 12 MP motion-capture cameras (Oqus 310 MRI, Qualisys, Göteborg, Sweden) were mounted in the scanner room, on the wall in front of the scanner entry, directly facing the scanner bed. In the optimal configuration to track the participants’ right hand in our scanner room, the cameras were distributed along the frontal wall of the scanner, in different positions along the horizontal and vertical axes. This configuration maximized the depth of the camera’s field of view to capture the inner part of the scanner bore where the participants’ hand is usually located. Three is estimated to be the minimum number of cameras to perform a 3D reconstruction of markers’ position in space, which in our case was sufficient to track the movements of the right forearm, which is the focus of our proof-of-concept.

Although the same cameras could theoretically track finger motion, the limited frontal field of view inside the scanner makes it difficult to reliably distinguish individual finger markers, resulting in multicollinearity across markers, affecting their identification. To address this, we included a separate device devoted explicitly to tracking finger flexion and extension. We used a kinematic glove designed for MR compatibility (Data Glove Ultra, Fifth Dimension Technologies, Pretoria, South Africa), equipped with 14 sensors (two for each finger, plus four between fingers). In the current implementation, we used a single glove designed for the right hand. It is important to note that the glove captures only finger flexion/extension while being uninformative about wrist or forearm position or orientation, which were instead obtained from the cameras.

The camera system and the glove were connected through shielded cables and interface devices to the computer in the control room via high-speed serial connections. We used a single high-end PC running Windows 11 as the control computer, running 3D kinematic modelling, data streaming and logging, and stereoscopic stimulus presentation. However, in different scenarios, these tasks could be easily assigned to different computers connected through the local network.

The key software components running on the control computer were Qualysis Track Manager (QTM), a vendor-provided Windows application allowing to capture data from the cameras, identify markers, and reconstruct a 3D representation of the captured motion, and Unity 6 (Unity Technologies, San Francisco, USA), which was used to stream data from QTM via TCP/IP and directly from the glove via serial connection, using open-source plugins provided by the hardware vendors, and to animate a 3D avatar accordingly.

Finally, the virtual environment produced by Unity was fed into high-definition MR-compatible goggles (VisualSystem HD, Nordic Neurolab, Bergen, Norway), featuring a 1920×1080 resolution at 60 Hz with a separate display for each eye, driven via a separate HDMI port. The device also supports eye tracking, although we have not yet incorporated this feature.

### 2.2 Procedure

#### 2.2.1 System calibration

At the beginning of each acquisition session, a dedicated calibration procedure was needed to let QTM software precisely locate the camera system within a fixed room-based coordinate system. This was done before the participant entered the scanner room and was required to place a reference object (a stationary L-shaped structure with four markers on it) at a predefined location in the scanner room (we positioned it at the top-left corner of the MRI table). Calibration consists of a 1-minute kinematic recording. An experimenter moves a T-shaped object of known size with three markers, waving it around where the participant’s hand will be located. In our experience, repeating the calibration procedure one to three times is often necessary until QTM produces a satisfying solution, consisting of a 3D view of the scanner room with the cameras located correctly.

Although the room coordinate system can be defined in several ways, we set it to correspond to the RAS (right-anterior-superior) standard which is ubiquitous in neuroimaging community, with the origin at the scanner isocenter, the Z axis aligned with the main body axis and pointing towards the head, the Y axis pointing up, and the X axis pointing to the right side of the participant body.

#### 2.2.2 Participant preparation

After entering the scanner room, the participant was asked to rest on the MRI table in a comfortable position. An experimenter assisted the participant in wearing the kinematic glove on his/her right hand. We placed a rubber glove on top of it to increase its stickiness to the skin. Then, we positioned five passive kinematic markers (diameter: 6.5 mm) on the participant’s right arm and hand (Figure 2A), following a standard layout defined by QTM (Animation Marker Set): 1) on the bony prominence of the outer elbow (”elbow-out”), 2) on the forearm at about half-distance between the elbow and the wrist (”forearm roll”), 3) on the outer (pinky side) and 4) the inner (thumb side) bones of the wrist (”wrist-out” and “wrist-in”), and 5) below the ring finger’s knuckle of the right hand (”hand-out”).

**Figure 2.**
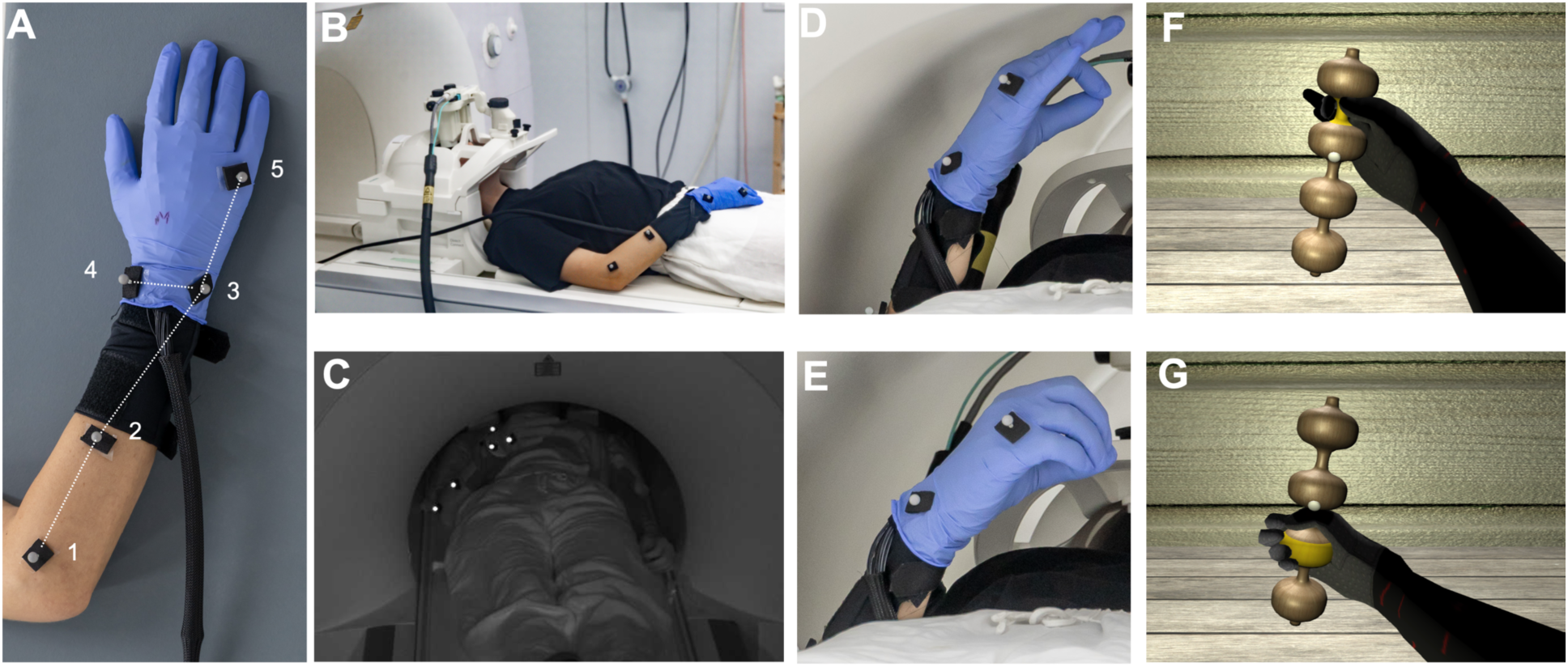
Detailed overview of MOTUM components. **A.** Location of markers on the participant’s arms: 1) elbow-out, 2) forearm roll, 3) wrist-out, 4) wrist-in, 5) hand-out (see main text for details). Dotted lines show the virtual bones reconstructed by the QTM software from the marker positions. The participant wears the kinematic glove, with a rubber glove on top of it to increase stickiness. **B.** Calibration of the kinematic acquisition was performed with the participant wearing the markers, the glove, the goggles, and the bed positioned outside the scanner. **C.** Example frame captured by the camera during the execution of a reaching-to-grasp movement within the scanner, where refractions from the markers are visible. **D-E.** Participant executing a precision (D) and a power (F) grip during the experiment. **F-G.** Visual feedback projected to the participant while executing the movements in D and E, respectively.

The participant was then prepared as usual for an fMRI session: headphones and soft cushions were placed within the base coil, and the upper coil was mounted on their head. We used a standard Siemens 64-channel head coil, which has enough space to mount and set up the VR goggles. The goggles were positioned on the top of the coil, and participants were guided in regulating the pupillary distance and the distance of the goggles from the eyes. The VR environment was initialized, and the goggles began receiving and displaying the VR scene.

Participants were then asked to perform four hand gestures, as cued by example images delivered to the goggles, to calibrate the kinematic glove by setting the maximum and minimum flexion values for each sensor. The four requested poses were: a) fist; b) fist with the thumb embraced by the other fingers; c) open hand; d) flat hand. The maximum and minimum flexion values were stored for successive loading.

#### 2.2.3 Marker identification

A key feature of the QTM system is the automatic identification of markers (AIM). With this function, QTM automatically identifies and labels movement trajectories, a crucial aid to maintaining a stable reproduction of the movement even when markers are not completely visible during the entire movement. AIM uses machine-learning models, which must be trained by multiple trials in which the specific motion to be tracked is recorded, in our case consisting of a set of repetitions of the reaching-to-grasp movement. It is crucial that the markers are correctly tracked throughout the whole model training phase. We experienced that this was hardly achievable when participants were entirely inside the scanner, as during the acquisition of MR images. The solution we found consists in acquiring a good amount of AIM trials (typically 1-2 minutes of recording where all markers were constantly visible) when the participant was lying on the MRI table but with the table set at about 35 cm from the magnet isocenter, so that the participant’s right hand was entirely outside the scanner bore (Figure 2B).

The first three volunteers trained the model more extensively by performing multiple reach-to-grasp movements with different starting positions and directions. An experimenter inside the scanning room checked the amplitude of the movement and that the markers were visible during the entire movement. Approximately 20 movements were captured in QTM, and this recording was then used to boost the automatic identification of trajectories. The following volunteers performed a shorter training phase. Note that we did not train a separate model for each participant: instead, the AIM process improves when new participants are added to the model, providing more examples of distances and angles between markers, i.e., depending on participants’ arm length. Therefore, each recording was added to the previously stored model to help the software apply it more easily for future participants.

The “skeleton solver” procedure in QTM allows for connecting the five markers through virtual bones describing the anatomy and biomechanical constraints of the forearm and hand (Figure 2A). Virtual bones formed “segments” whose 3D position and orientation in room coordinates are streamed in real-time at 120 Hz by QTM to the Unity software.

#### 2.2.4 Environment calibration

Once the AIM model was trained and skeleton data was correctly streamed, the scanner table was moved to the final position, reaching the magnet isocenter.

Participants were asked to engage in the grasping movements they had been instructed to perform. Importantly, volunteers were instructed not to move their head during the reach-to-grasp action and to keep the right shoulder relaxed during the movement. Kinematic skeleton data fed into Unity was used to animate a humanoid avatar through custom scripts. In this way, participants could see their hand movements reproduced on the avatar in a first-person perspective. A frame-wise interpolation was applied to the movement data to compensate for occasional marker loss, using a smoothing window of 5 frames.

Using a customized graphical user interface (GUI), the experimenter could modify the position of several components of the virtual environment, tailoring them to match the individual range of motion of the participant, e.g., adapting to different arm lengths. In this phase, it was crucial to ensure that the movement of interest occurred within the camera’s field of view, where marker positions could be accurately detected. The GUI also allowed control of the size and position of the grasping target and several camera settings, such as 3D distortions, eye convergence, field of view (FOV), and pupillary distance.

The participants’ right hand was positioned over their abdomen as the starting position for the movement. In the virtual environment, participants saw their right forearm and hand resting on a table as if they were sitting on a chair in front of the table. Although the real and virtual positions of the body and hand conflicted, the visual experience was immersive enough to make participants immediately feel the virtual hand as their own. We let subjects move their hand and fingers for a couple of minutes while observing the virtual counterpart of their movement, to increase this ownership feeling. In the setup described here, the reach-to-grasp action had to be executed by extending the arm and moving the hand towards the feet. In the virtual world, this corresponded to moving the arm and hand forward to grasp an object on the virtual table.

### 2.3 Participants

Eight volunteers participated in this proof-of-concept study. One participant was excluded due to artefacts caused by hair gel, so the final sample was composed of seven participants (5 females; age: 26.86 ± 4.38). To avoid the scanner bore obstructing the view of the cameras, we excluded participants shorter than 155 cm. Participants were right-handed, as assessed by the Edinburgh Handedness Inventory (Oldfield, 1971), and had normal or corrected-to-normal vision. Participants gave their written informed consent to participate in the study. The study was approved by the regional research ethics committee (Prot. CE/PROG.932).

### 2.4 Stimuli and task

A VR environment was developed in Unity software editor (version 6.0). The scene was composed of a wooden background wall, a wooden table, and on top of it, a 3D target object. The latter was crafted through coding to generate a symmetric sinusoidal shape with four alternating wide and narrow sections, which could be grasped through a power and a precision grip, respectively (Figure 2F-G). The object was tilted at 20° on the Z-axis to facilitate grasping movements on its whole length. A white sphere in the object’s center acted as a fixation point. The object and the table were positioned in front of a humanoid avatar, of which only the arm was visible to the participants.

Participants were instructed to start each trial with their right palm positioned on their abdomen, in a comfortable position, and to fixate the fixation point throughout the entire acquisition. At the beginning of each trial, a yellow light highlighted one of the four central sections of the virtual object for 2.5 s. Notably, the light could appear on a narrow or a wide section of the object, instructing a precision (“pinching” with thumb and index finger) or a power grip (closing the whole hand), respectively. Trials were organized in blocks of three, preceded by an auditory instruction (1s) prompting participants to “move” or “observe”. During “Movement” blocks, participants had to reach the light target with the hand while performing either a precision or a power grip, and then immediately return to the starting position. During “No Movement” blocks (NM), participants observed the scene, including the light appearing on the targeted section, but performed no movements.

To disentangle the effect of the visual feedback, in half of the “Movement” blocks (“Move Visible” blocks; MV), participants were given complete visual feedback of their movement, with the virtual hand moving based on the real-time streaming of the participant’s hand movements. In the other half of “Movement” blocks (“Move Invisible” blocks; MI), participants had no visual feedback, with the virtual hand remaining in the starting position irrespective of the movements performed by the participant.

Trials were spaced by a jittered interval varying according to an exponential distribution from 5 to 13 s (mean: 7 s). Blocks were interleaved following a pseudo-random sequence. Each of the three block types occurred five times in each of the four total functional scans acquired per participant, for a total of 60 trials per condition.

After the scan, participants were asked to complete a six-item questionnaire to assess the degree of embodiment on a 10-point rating scale (Tieri et al., 2015; Botvinick & Cohen, 1998). Two measures are obtained: 1) feeling of ownership, i.e., the degree to which the participants felt that the virtual arm belonged to them (average of responses to Items 1-2; Item 3 acted as a control); 2) agency, i.e., the feeling to be in control of the virtual hand’s action (average of responses to Items 4-5; Item 6 acted as a control).

### 2.5 Image acquisition

For each participant, we acquired:

1. a T1-weighted structural image with a magnetization-prepared rapid gradient-echo (MPRAGE) sequence with perspective motion correction and selective reacquisition of data corrupted by motion based on interleaved 3-D EPI navigators (Tisdall et al., 2012; Hess et al., 2011). Volumetric imaging included 176 slices, isotropic resolution = 1 mm^3^, TR = 2500 msec, TE = 1.67 msec, inversion time = 1080 msec, flip angle = 8°;
2. a T2-weighted structural image using a 3D SPACE sequence, which employs variable flip angle refocusing pulses with a nominal flip angle of 120°. The acquisition consisted of 1 mm isotropic resolution volumes, TR = 3200 msec, TE = 564 msec. The matrix size was 256 × 240, with a field of view of 240 × 225 mm;
3. five functional, whole-brain MR images acquired with a T2٭-weighted gradient-echo EPI sequence, a multiband factor of 6, and an isotropic voxel size of 2.4 mm^3^ (60 slices, field of view = 88 × 88 mm^2^, TR = 0.800 s, TE = 0.03 s, flip angle = 52°, no in-plane acceleration; Xu et al., 2013; Feinberg et al., 2010; Moeller et al., 2010);
4. two spin-echo EPI volumes with phase encoding in opposite directions, no multiband acceleration, and the same geometrical and sampling properties of functional runs, acquired for field mapping (TE = 80 msec, TR = 7060 msec).

Participants underwent four task-based functional runs of variable length, depending on the individual trial list (mean volumes per run: 569.5; sd: 19.50), and one resting state run of 450 volumes. One participant did not complete the resting state run. For the current work, the resting state runs were only used to compare the amount of head movement with the runs requiring hand movements.

The experimental session lasted roughly 1h20m, involving setup, structural, and functional image acquisition.

### 2.6 Kinematic analyses

Kinematic data from arm and hand movements reconstructed in real time and streamed in the VR environment were saved from Unity at the experiment frame rate (typically 60 fps) and analyzed offline. Data included the three-dimensional position and rotation of the elbow, forearm, wrist, thumb tip, and index finger tip.

We first computed instantaneous arm velocity as the 3D Euclidean distance between wrist positions in consecutive frames divided by the corresponding time interval, smoothed using a 5-frame moving average window. We then applied a k-means clustering algorithm on smoothed arm velocity for each trial to segment periods where the hand was moving or was still. This allowed us to identify, for each trial: 1) the reaction time, defined as the moment in which the hand started to move (i.e., the first transition from a still to a moving period); 2) the reaching phase (i.e., the period from the reaction time to the next transition to a still period); 3) the grip phase (i.e., the still period in which the hand was close to the target); 4) and the back-movement phase, whose last time point was considered as the end of the trial.

Position and velocity data from each trial were plotted and visually inspected to identify anomalies in the participant’s behavior and/or kinematics segmentation, which were manually marked as error trials. After excluding such trials, we computed kinematic parameters separately for the reaching, grip, and back-movement phases. We quantify the movement in the reaching and back-movement phases by computing the total duration of the movement, the total distance traveled by the arm (based on the reconstructed wrist position), the maximum and average velocity, and two trajectory indices: (a) a trajectory curvature index, measured as the ratio of the actual path length to the straight-line distance between the starting and ending points; and (b) the maximum deviation, measured as the maximum perpendicular distance from the optimal straight-line path. The same parameters were also computed for the hand (finger) movements, based on the reconstructed position of the tip of the index finger relative to the wrist. Finally, to quantify kinematic parameters of the grip phase, we computed grip aperture as the Euclidean distance between the tip of the index finger and the thumb, and hand aperture as the Euclidean distance between the tip of the index finger and the wrist. We extracted the minimum, maximum, and average grip and hand aperture during the grip phase as meaningful kinematic parameters for further analyses.

In sum, we computed 28 kinematic parameters from each movement trial classified as correct. A principal component analysis (PCA) with varimax rotation was performed on the whole set of kinematic parameters across all subjects using Jamovi to identify a reduced set of independent kinematic components explaining most of the variance. The scores of each component were then used in the neuroimaging analyses to reveal a possible correlation between kinematics and brain activity (see below).

Finally, to quantify the impact of arm and hand movement on head movement and potential artefacts induced by arm and hand movement in fMRI data, we derived an estimate of the amount of arm and hand movement within each fMR time frame, by resampling the position of the wrist and of the tip of index finger at each TR (0.8s) and computing the framewise movement of the arm (based on the position of the wrist relative to the elbow) and of the hand (based on the position of the tip of the index relative to the wrist), using the same approach which is commonly used in fMRI for estimating the framewise movement of the head, i.e., the framewise displacement (FD) index based on the formula in Power et al. (2012). We adapted the original formula using a reference distance of 300 mm for forearm movement estimation (i.e., the average distance between the wrist and the elbow) and 180 mm for hand movement estimation (i.e., the average distance between the tip of the index and the wrist).

### 2.7 Image preprocessing

Results included in this manuscript come from preprocessing performed using fMRIPrep 25.0.0 (Esteban et al., 2019; Esteban et al., 2018), which is based on Nipype 1.9.2 (Gorgolewski et al., 2011; K. J. Gorgolewski et al., 2018). The following text was automatically generated by fMRIPrep with the express intention to be copied and pasted into manuscripts unchanged, and is released under the CC0 license.

#### 2.7.1 Anatomical data preprocessing

The T1w image was corrected for intensity non-uniformity (INU) with N4BiasFieldCorrection (Tustison et al., 2010), distributed with ANTs 2.5.4 (Avants et al., 2008), and used as T1w-reference throughout the workflow. The T1w-reference was then skull-stripped with a Nipype implementation of the antsBrainExtraction.sh workflow (from ANTs), using OASIS30ANTs as target template. Brain tissue segmentation of cerebrospinal fluid (CSF), white matter (WM), and gray matter (GM) was performed on the brain-extracted T1w using fast (FSL: Zhang, Brady, and Smith 2001). Brain surfaces were reconstructed using recon-all (FreeSurfer 7.3.2: Dale, Fischl, and Sereno 1999), and the brain mask estimated previously was refined with a custom variation of the method to reconcile ANTs-derived and FreeSurfer-derived segmentations of the cortical gray matter of Mindboggle (Klein et al., 2017). The T2-weighted image was used to improve pial surface refinement. Brain surfaces were reconstructed using recon-all (FreeSurfer 7.3.2: Dale, Fischl, and Sereno 1999), and the brain mask estimated previously was refined with a custom variation of the method to reconcile ANTs-derived and FreeSurfer-derived segmentations of the cortical gray matter of Mindboggle (Klein et al., 2017). Volume-based spatial normalization to two standard spaces (MNI152NLin6Asym, MNI152NLin2009cAsym) was performed through nonlinear registration with antsRegistration (ANTs 2.5.4), using brain-extracted versions of both T1w reference and the T1w template. The following templates were selected for spatial normalization and accessed with TemplateFlow (24.2.2, Ciric et al. 2022): FSL’s MNI ICBM 152 non-linear 6th Generation Asymmetric Average Brain Stereotaxic Registration Model [Evans et al. (2012); TemplateFlow ID: MNI152NLin6Asym], ICBM 152 Nonlinear Asymmetrical template version 2009c [Fonov et al. (2009); TemplateFlow ID: MNI152NLin2009cAsym]. Grayordinate “dscalar” files containing 91k samples were resampled onto fsLR using the Connectome Workbench (Glasser et al., 2013).

#### 2.7.2 Functional data preprocessing

For each of the BOLD runs, the following preprocessing was performed. First, a reference volume was generated using a custom methodology of fMRIPrep for head motion correction. Head-motion parameters with respect to the BOLD reference (transformation matrices and six corresponding rotation and translation parameters) are estimated before any spatiotemporal filtering using mcflirt (FSL, Jenkinson et al. 2002). The estimated fieldmap was then aligned with rigid registration to the target EPI (echo-planar imaging) reference run. The field coefficients were mapped onto the reference EPI using the transform. The BOLD reference was then co-registered to the T1w reference using bbregister (FreeSurfer), which implements boundary-based registration (Greve & Fischl, 2009). Co-registration was configured with six degrees of freedom. The aligned T2w image was used for initial co-registration. Several confounding time series were calculated based on the preprocessed BOLD: framewise displacement (FD), DVARS, and three region-wise global signals. FD was computed using two formulations following Power (absolute sum of relative motions, Power et al. (2014)) and Jenkinson (relative root mean square displacement between affines, Jenkinson et al. (2002)). FD and DVARS are calculated for each functional run, both using their implementations in Nipype (following the definitions by Power et al. 2014). The three global signals are extracted within the CSF, the WM, and the whole-brain masks. Additionally, physiological regressors were extracted to allow for component-based noise correction (CompCor, Behzadi et al. 2007). Principal components are estimated after high-pass filtering the preprocessed BOLD time-series (using a discrete cosine filter with 128s cut-off) for the two CompCor variants: temporal (tCompCor) and anatomical (aCompCor). tCompCor components are then calculated from the top 2% variable voxels within the brain mask. For aCompCor, three probabilistic masks (CSF, WM and combined CSF+WM) are generated in anatomical space. The implementation differs from that of Behzadi et al. in that instead of eroding the masks by 2 pixels on BOLD space, a mask of pixels that likely contain a volume fraction of GM is subtracted from the aCompCor masks. This mask is obtained by dilating a GM mask extracted from the FreeSurfer’s aseg segmentation, and it ensures components are not extracted from voxels containing a minimal fraction of GM. Finally, these masks are resampled into BOLD space and binarized by thresholding at 0.99 (as in the original implementation). Components are also calculated separately within the WM and CSF masks. For each CompCor decomposition, the k components with the largest singular values are retained, such that the retained components’ time series are sufficient to explain 50 percent of the variance across the nuisance mask (CSF, WM, combined, or temporal). The remaining components are dropped from consideration. The head-motion estimates calculated in the correction step were also placed within the corresponding confounds file. The confound time series derived from head motion estimates and global signals were expanded with the inclusion of temporal derivatives and quadratic terms for each (Satterthwaite et al., 2013). Frames that exceeded a threshold of 0.5 mm FD or 1.5 standardized DVARS were annotated as motion outliers. Additional nuisance timeseries are calculated by means of principal components analysis of the signal found within a thin band (crown) of voxels around the edge of the brain, as proposed by Patriat, Reynolds, and Birn (2017). The BOLD time series were resampled onto the left/right-symmetric template “fsLR” using the Connectome Workbench (Glasser et al., 2013). A “goodvoxels” mask was applied during volume-to-surface sampling in fsLR space, excluding voxels whose time series have a locally high coefficient of variation. Grayordinates files (Glasser et al., 2013) containing 91k samples were also generated with surface data transformed directly to fsLR space and subcortical data transformed to 2 mm resolution MNI152NLin6Asym space. All resamplings can be performed with a single interpolation step by composing all the pertinent transformations (i.e., head-motion transform matrices, susceptibility distortion correction when available, and co-registrations to anatomical and output spaces). Gridded (volumetric) resamplings were performed using nitransforms, configured with cubic B-spline interpolation.

Finally, a spatial smoothing was performed on the cortical surface using geodesic smoothing with a 6 mm full-width at half-maximum (FWHM) kernel (Hagler et al., 2006).

### 2.8 Image analysis

We analyzed functional images for each participant in the surface-based fsLR space on a vertex-by-vertex basis, implementing a general linear model (GLM) in SPM12. Each trial was modeled as a canonical hemodynamic response function time-locked to the trial onset. Separate predictors were entered for the three trial types: “Move Visible”, “Move Invisible”, and “No Movement”. Error trials with and without participant movements were modeled as separate regressors. The model included framewise displacement and five CompCor components as a standard way to control for head movement-related artifacts. We also included frame-wise estimates of arm and hand movement derived from kinematic analyses (see above) to control for possible artifacts induced by arm and hand, in addition to head, movements, after orthogonalization relative to head frame-wise displacement and orthogonalization of hand relative to arm frame-wise displacement.

As a proof-of-concept for the present study, we included in the general linear model, as linear modulators of the activity of each “Move Visible” and “Move Invisible” trial, the principal component scores extracted from the PCA analysis of kinematic indices (see above), to show whether the variance in kinematic aspects of reach-to-grasp movements is associated to specific patterns of brain activity. This approach aims to show whether MOTUM allows not only to perform real-time motion tracking in fMRI experiments, but also to uncover relationships between brain activity and movement kinematics.

Individual statistical parametric maps were obtained for the comparison of the three conditions against each other and the overall effect of the kinematic indices derived from the PCA. Statistical maps were thresholded at *p* < 0.05 corrected for multiple comparisons at the cluster level through a topological false discovery rate procedure based on random field theory (Chumbley et al., 2010), after applying a cluster-forming threshold of *p* < 0.001 uncorrected.

## 3. Results

### 3.1 Quality of the virtual reality experience

Providing a good-quality immersive virtual reality experience to participants in an environment subject to strict physical and technical constraints, such as MR, is a difficult challenge. One crucial aspect is the possible mismatch between the proposed virtual environment and the real environment in which the experience is conducted. Our choice was to create a virtual world reproducing a typical situation of everyday life (sitting in front of a table), although, of course, it might create a mismatch with the real position of the subject (supine on the MR bed). However, participants reported their experience as highly natural and reported no difficulty adapting to and feeling immersed in the virtual environment.

A second crucial aspect to evaluate the quality of the virtual reality experience is how well subjects’ movements are reflected in the real-time sensory feedback given to them. In our setup, feedback was limited to the visual modality, and, unlike in commonly available VR setups based on head-mounted displays, it was not possible to move one’s head and explore the environment from different viewpoints, given the inherent limitations of the MR setup. Critically, however, the virtual world included a real-time avatar moving in sync with the real subject’s hand and arm. The real-time reconstruction was sufficiently accurate and fast, as reflected by participants’ reports of feeling of ownership of the avatar’s arm and hand (average scores to Items 1-2 of the Embodiment questionnaire: “I felt as if I was looking at my hand” and “I felt that the virtual hand was my own”; 5.57, s.d. 2.5) and a good sense of agency of the observed arm/hand movements (average scores to Items 4-5 of the Embodiment questionnaire: “I perceived the movements of the virtual hand as if they were my movements” and “I felt that I had caused one or more movements of the virtual hand”; 6.92, s.d. 2.5).

In conclusion, given the inherent constraints of the MR environment, we succeeded in providing an immersive experience of good quality, generating a satisfying level of embodiment and sense of agency over the virtual avatar.

### 3.2 Quality of arm, hand, and finger tracking

The system successfully tracked and streamed the reach-to-grasp kinematics of all participants in real-time: loss of tracking of arm and hand markers, probably due to physical occlusion, was extremely rare, and compromised only 7 out of 1260 total trials. Finger flexion and extension obtained through the glove were consistently acquired and streamed with no issues.

A bug in a preliminary version of the Unity stimulus presentation packages resulted in incorrect visual feedback given to participants despite correct marker tracking, compromising four sets of consecutive trials in different runs (43 trials in total). For all other trials, we visually inspected position, velocity, and grip/hand aperture plots and identified no visible anomalies in marker tracking and velocity profiles. In 9 cases, participants performed unexpected hand movements during a movement trial (e.g., they repeated the reaching movement twice), while in 17 further cases, they moved the arm in a no-movement trial.

We also visually inspected the results of the automated identification of reaching, grip, and back-movement phases for all remaining movement trials. In 31 out of 840 movement trials (3.7%), the algorithm failed in correctly parsing the three movement phases. The final set of accepted trials included 377 “Move Visible”, 388 “Move Invisible”, and 388 “No Movement” trials. All problematic trials were excluded from the kinematic analyses and marked as error trials in the fMRI analysis.

### 3.3 Quality of fMRI recordings during arm/hand movements

To determine whether participants’ arm and hand movements significantly affected the quality of fMRI recordings, we performed several analyses on quality control measures.

#### 3.3.1 Head movement induced by arm/hand movements

First, we assessed whether moving the arm and hand during reaching-to-grasp trials induced excessive head movement, compromising the quality of the recordings. We quantified head movement using the framewise displacement index (FD), which is considered a standard measure of instantaneous head movement occurring at each MR frame. Supplementary Figure 1 shows the FD time series for each run across all subjects. Figure 3 (left panel) shows the spread and density of FD values during movement and no-movement blocks in the main fMRI runs, and, for the sake of comparison, during resting-state runs. Mean FD values were 0.0961 mm (s.d. 0.0463) during movement blocks, 0.0853 mm (s.d. 0.0813) during no-movement blocks, and 0.0801 mm (s.d. 0.0525) during resting-state scans. As expected, moving the arm and hand increased the amount of head movement relative to no-movement blocks (t_6_ = 3.4454, *p* = 0.0137), while the difference between movement blocks and resting-state scans was close to significance (t_5_ = 2.5061, *p* = 0.0541).

**Figure 3.**
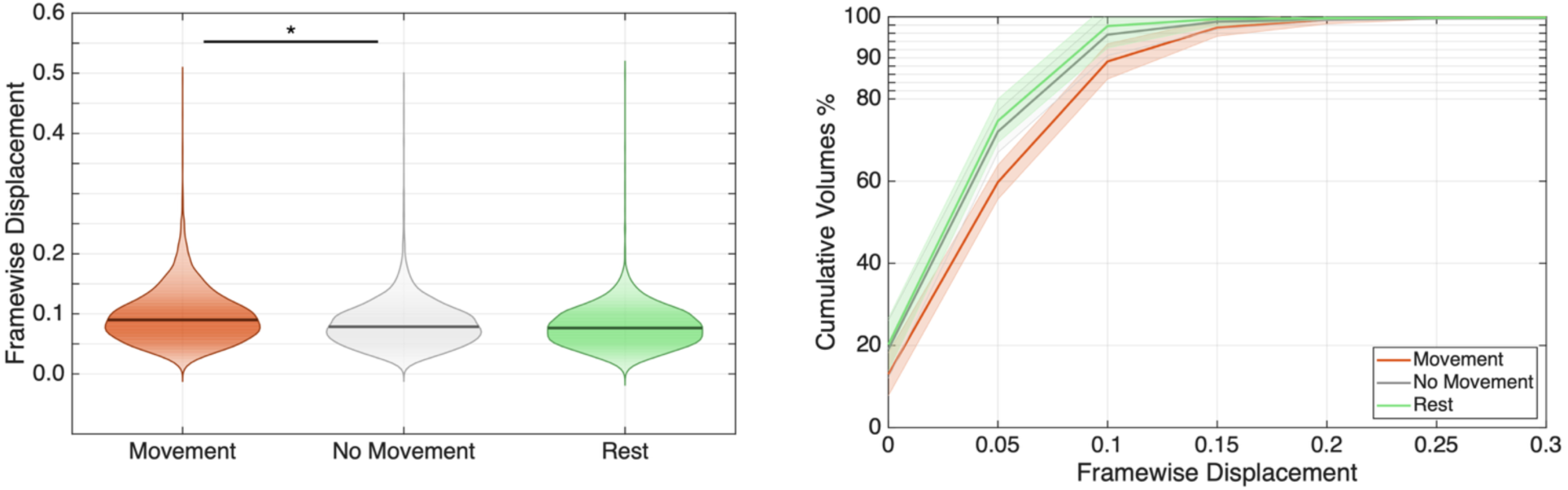
Assessment of head movements during reach-to-grasp execution. **Left panel:** violin plots of framewise displacement (FD) distribution across conditions. Outliers above 0.5 mm were excluded for better visualization. The violin shapes show the probability density of FD values, with wider sections indicating more frequent values. The horizontal black line within each violin represents the median FD value. \**p* < 0.05. **Right panel**: cumulative distribution of framewise displacement (FD) across conditions showing the progressive proportion of volumes retained at different FD thresholds. The x-axis represents FD values (mm), and the y-axis shows the cumulative percentage of volumes with FD below each threshold value. Shaded areas indicate standard error across subjects. For both panels, colors for each condition are as follows: Movement = orange; No Movement = grey; Rest = green.

However, FD values consistently remained within more than acceptable limits for task-based fMRI studies: 99.9% of MR frames during movement blocks (as well as 99.8% during no-movement blocks and 99.9% during resting-state scans) fell below the commonly used 0.5 mm threshold for FD (Siegel et al., 2014). Even considering a more stringent 0.2 mm threshold, 97.4% of volumes during movement blocks were retained (98.8% during no-movement blocks and 99.5% during resting-state scans). Fig. 3 (right) shows the simulation of a volume censoring approach across multiple FD thresholds, demonstrating that even at stringent thresholds, all conditions maintained adequate temporal coverage (> 5.9 minutes of retained data for each condition for each subject), confirming the feasibility of motion-sensitive preprocessing approaches while preserving statistical power.

A more direct way to show that arm/hand movements induce head movement consists of the frame-by-frame correlation between indicators of head movement, such as FD, and the corresponding indicators of frame-by-frame arm and hand (finger) movements, computed through an FD-like approach (see Methods for details). The correlation of FD with hand framewise movement, calculated during movement blocks, was moderately positive (arm: r = 0.229, s.d. 0.08; hand: r = 0.199, s.d. 0.03); both were significantly higher than zero (Fisher’s transform: hand: t_6_ = 7.30, *p* < 0.001; index: t_6_ = 16.77, *p* < 0.001).

Root mean square displacement (RMSD) is another head movement-related index representing the head’s cumulative displacement over the entire scanning session, providing an overall measure of head stability. RMSD values were 0.053 mm (SD = 0.026) for movement blocks, 0.047 mm (SD = 0.045) for no-movement blocks, and 0.045 mm (SD = 0.030) for resting-state scans. Despite a slight increase in movement compared to no-movement blocks (t_6_ = 3.50, *p* = 0.0128), RMSD remained well within acceptable limits (< 0.1 mm) across all conditions.

In conclusion, the reaching-to-grasp movement caused a noticeable but slight increase in head movement during the scan, which remained well below the guard levels.

#### 3.3.2 Signal artifacts induced by arm/hand movements

To verify whether arm and hand movements were responsible for signal artifacts in fMR images not necessarily related to head movement, we considered standardized DVARS (temporal derivative of RMS variance over voxels), which is a standard index which quantifies the rate of change of BOLD signal intensity across the whole brain from volume to volume, with higher values indicating greater signal fluctuation potentially related to motion artifacts. Standardized DVARS provides a normalized measure that accounts for baseline signal variance, making comparisons across subjects and sessions reliable. Mean standardized DVARS values were 1.16 (s.d. = 0.15) for movement blocks, 1.14 (s.d. 0.15) for no-movement blocks, and 1.19 (s.d. 0.14) for resting-state scans. As in the case of framewise displacement, a noticeable but minimal increase for movement blocks could be observed, (t_6_ = 3.72, *p* = 0.01), although the frame-by-frame signal variability remained well below acceptable limits (standardized DVARS < 1.5: Esteban et al., 2019; but see also Afyouni & Nichols, 2018) and not significantly different from the levels of resting-state scans on this sample (t_5_ = 0.66, *p* = 0.53).

A more direct way to assess and control for the artifactual effects induced by arm/hand movements is to include a measure of frame-by-frame movement as a predictor in the general linear model. We derived two measures similar to framewise displacement, quantifying the amount of arm and hand (finger) movements in each frame, respectively (see Methods for details), and included them in the general linear models (Figure 4). Results revealed that in 6 out of 7 participants, hand (finger) movements showed little or no correlation with brain activity; in only one participant (S4), we observed hand-movement-related artefacts in frontal and parietal reaching areas in both hemispheres. Instead, arm movements induced a bigger effect on the recorded brain activity, with an increased displacement correlated with heightened activation of temporal, parietal, and frontal areas in 5 out of 7 participants. In conclusion, we found evidence of a direct artifactual effect of only arm movements on the BOLD signal, which, however, was quantitatively small and controlled for in the GLM design.

**Figure 4.**
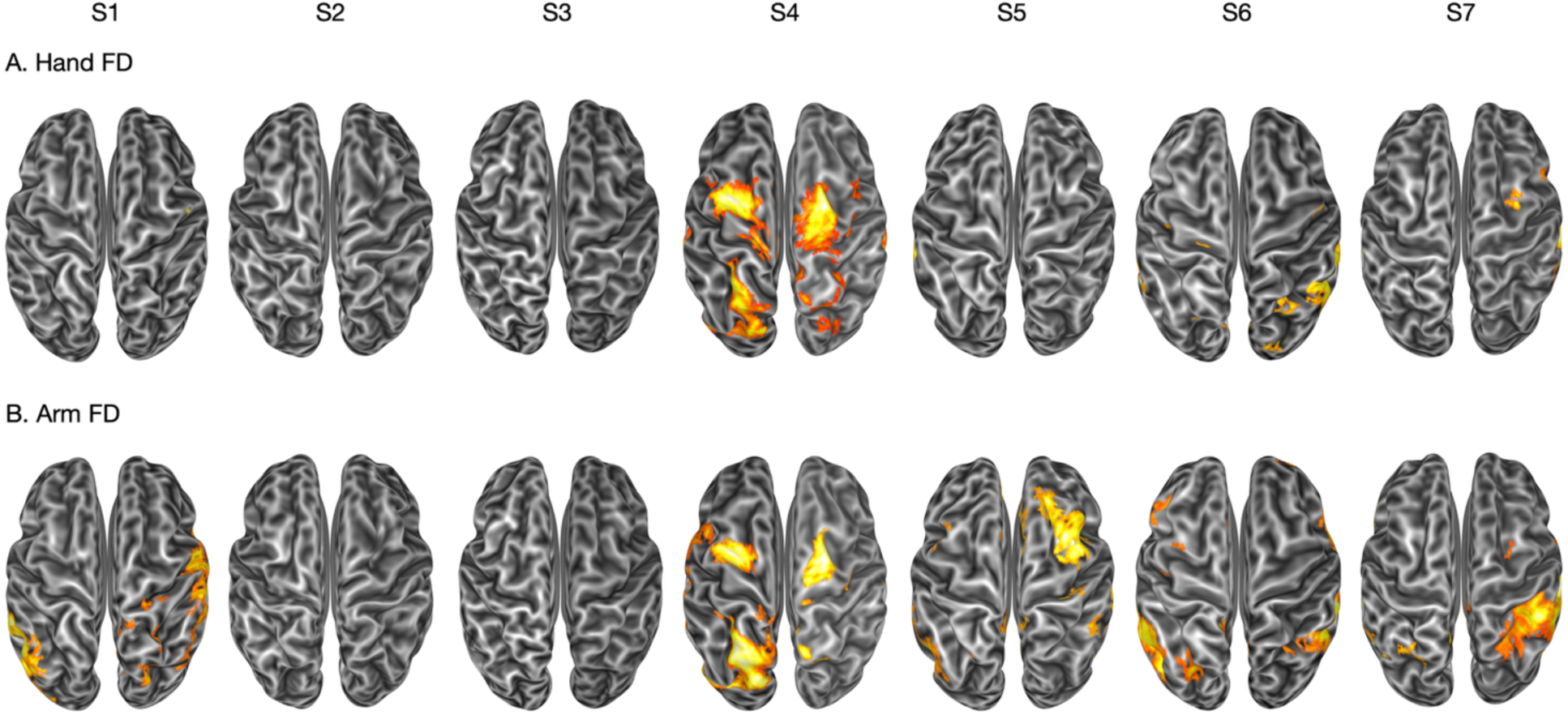
Effect of the hand and arm movements on brain activity. For each participant, an inflated dorsal view of their surface reconstruction is shown, with regions showing a significant (p < 0.05 corrected for false discovery rate at the cluster level, with a cluster-forming threshold of p < 0.001 uncorrected) effect of the hand (**A**) and arm (**B**) movements on the brain activation.

### 3.4 Kinematic results

Figure 5 shows an example of kinematic data for a typical reaching trial, showing the temporal profile of arm velocity and grip aperture and the automatic parsing of reaching, grip, and back-movement phases.

**Figure 5.**
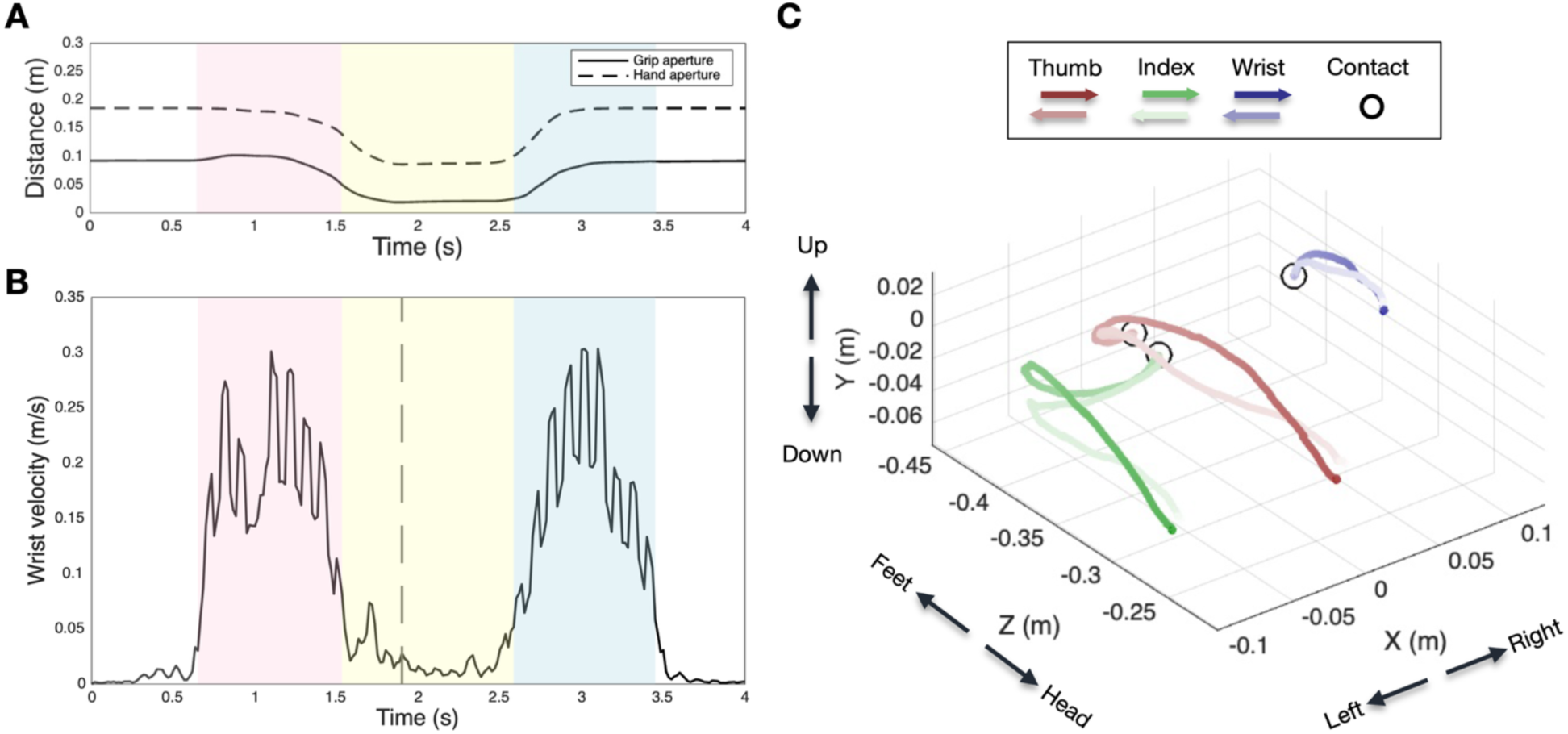
An example of kinematics from a single trial of an individual participant**. Left panel, top:** index-to-thumb (grip aperture; solid line) and index-to-wrist (hand aperture; dashed line) Euclidean distances across time. **Left panel, bottom:** wrist velocity profile across time. Color bands indicate phase segmentation, as computed by a classification algorithm on wrist velocity profiles: pink = reaching phase; yellow = grip phase; blue = back movement phase. **Right panel:** 3D positions of thumb (red), index (green), and wrist (blue) throughout a trial. The moment of the trajectory in which the participant virtually touched the target is indicated with a black circle.

The reaching movement started on average 624 ms after the visual cue onset (average across seven participants; s.d. 85 ms) and lasted on average 789 ms (s.d., 166 ms). The grip phase lasted on average 1132 ms (s.d. 625 ms), while the back-movement phase lasted 1040 ms (s.d. 232 ms). Grip aperture measures showed significant differences between power and precision grips. The minimum grip aperture was 45 mm (across-subject average; s.d. 6.7) when performing a power grip and 72 mm (s.d. 14) when performing a precision grip (t_6_ = 5.44, *p* = 0.002). Hand grip measures showed larger differences, with a minimum aperture of 122 mm (s.d. 28) when performing a power grip versus 162 mm (s.d. 12) when performing a precision grip (t_6_ = 4.244, *p* = 0.005). This difference reflects biomechanical properties of each grip type, where wrist-to-index distance is maximal with open hand, intermediate during precision grip, shorter during power grip, and minimal with closed hand (Figure 2D-E). In precision grip, the index finger flexes primarily at the metacarpophalangeal joint (knuckle) while the distal and proximal interphalangeal joints remain relatively extended, optimizing fine motor control. In contrast, power grip involves progressive flexion of all finger joints as the entire hand closes around the object, creating maximum contact surface and grip strength through distributed force across multiple joints.

The principal component analysis conducted on a set of 28 kinematic measures extracted from each trial (including distance traveled, maximum and average velocity, curvature index, and maximum deviation of the wrist and the index trajectory in the reaching and back-movement phases, duration of the reaching and back-movement phases, and minimum, maximum, and average grip and hand aperture during the grip phase) identified seven factors based on parallel analysis, explaining 7-13% of total variance each, and 75% in total. Interestingly, these factors explained in a partially independent way the variance of the reaching and back-movement phases and the grip, with independent components representing the arm/hand velocity and the distance traveled in the two phases, the grip/hand aperture, and the trajectory curvature (see Table 1). Component loadings are shown in Supplementary Table 1.

**Table 1.** Principal components extracted from kinematic analysis.

| Component | Variance % | Name |
| --- | --- | --- |
| 1 | 13.19 | Arm velocity (reaching and back-movement phases) |
| 2 | 12.36 | Distance traveled by arm and hand (reaching phase) |
| 3 | 9.78 | Distance traveled by arm and hand (back-movement phase) |
| 4 | 9.67 | Arm and hand velocity (reaching phase) |
| 5 | 10.49 | Arm and hand velocity (back-movement phase) |
| 6 | 11.89 | Grip and hand aperture |
| 7 | 7.41 | Trajectory curvature |
| Total | 74.79 |  |

### 3.5 Brain activation induced by reaching-to-grasp movements

The whole-brain trial-design GLM results are shown in Figure 6 on the surface reconstructions of each of the seven volunteers.

**Figure 6.**
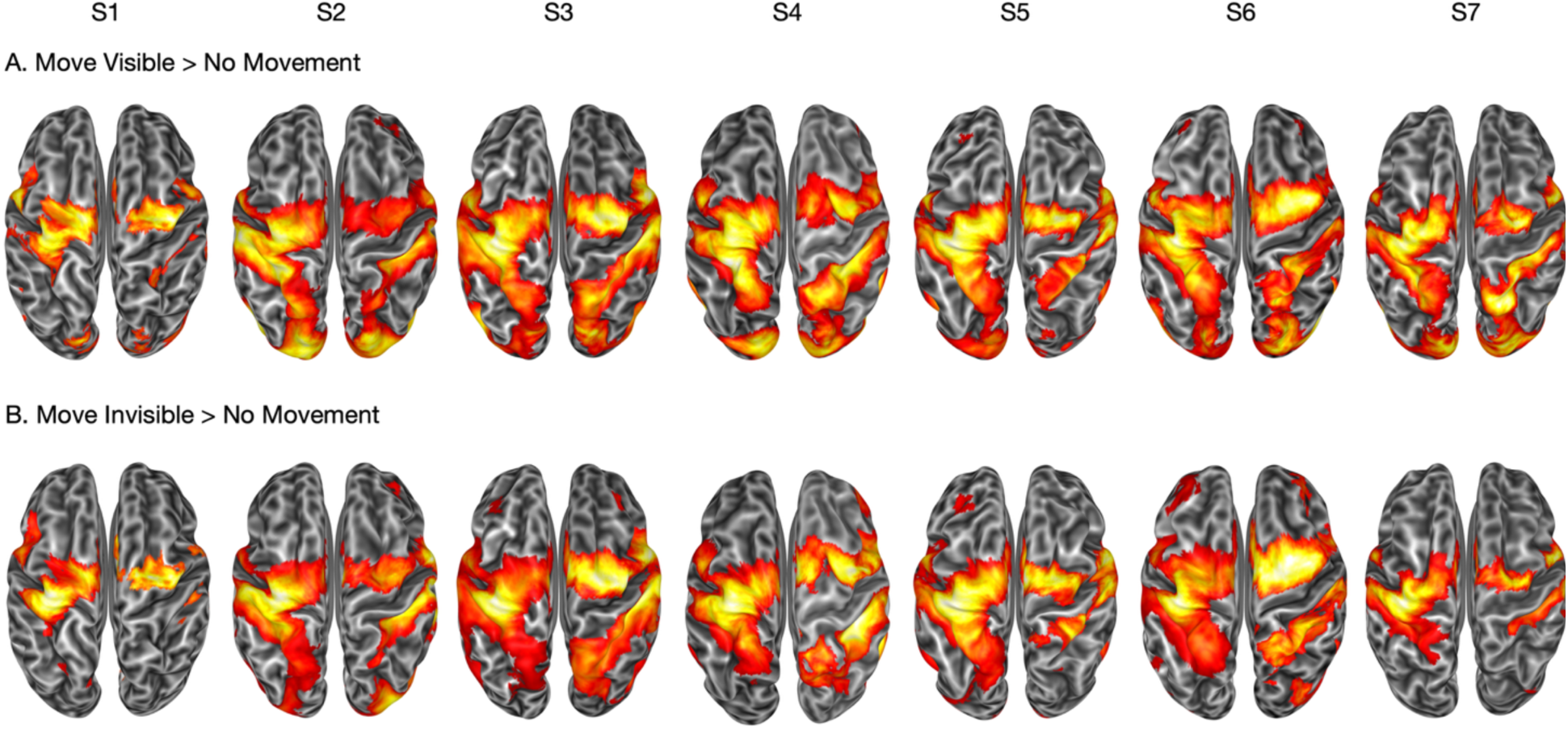
Individual brain activation maps for 7 participants during the two movement conditions. Top row: Regions with significantly greater activation during reach-to-grasp movements with visual feedback of the moving hand (Move Visible > No Movement). Bottom row: Regions with significantly greater activation during reach-to-grasp movements without visual feedback (Move Visible > No Movement). Each column represents individual-level analysis from one participant. Statistical parametric maps are displayed on the individual surface reconstruction, thresholded at p < 0.001 (uncorrected).

When compared to static trials (i.e., NM trials), movement execution devoid of visual feedback increased activity in bilateral parietal reach- and grasp-related areas (MIP, AIP), extending to the supramarginal gyrus, and in the pre-SMA (*p* <0.001). Only in the left hemisphere, contralateral to the moving hand, we detected activity of somatomotor areas encompassing PMd, M1, and S1, and extending to the medial bank (SMA) (Figure 6, top row). Performing a reach-to-grasp movement with visual feedback additionally activated bilateral occipital and visual processing regions, including area MT+ and the intraoccipital sulcus, as well as SPL, IPS, and PMd, bilaterally (Figure 6, bottom row). This is expected, as visual areas (i.e., MT+, intraoccipital sulcus) likely transmit visuo-spatial information to parietal reaching system regions (SPL), then to frontal motor (PMd) areas through the superior longitudinal fasciculus I (SLF I), creating a visuomotor control loop that enables adaptive movement within the novel virtual context.

### 3.6 Modulation of brain activity induced by kinematic parameters of reaching-to-grasp movements

The conjunction analysis of all seven PCA components revealed widespread significant modulations across distributed brain regions (Figure 7), with substantial interindividual differences observed across parieto-frontal networks and crucial contributions from occipital areas. Taken together, key regions showing significant parametric modulation included the angular gyrus (AG), the posterior intraparietal sulcus (pIPS), the primary somatosensory cortex (S1), the middle frontal gyrus (MFG), area V6, and MT+. Although the small sample of the current study prevents strong statistical analyses, a tendency of enhanced modulation by kinematic indices on the activity of posterior parietal and frontal regions during the “Move Invisible” condition compared to the “Move Visible” condition emerges. This may be explained by the fact that the MI condition relies more heavily on effector-based control mechanisms than visually guided movement.

**Figure 7.**
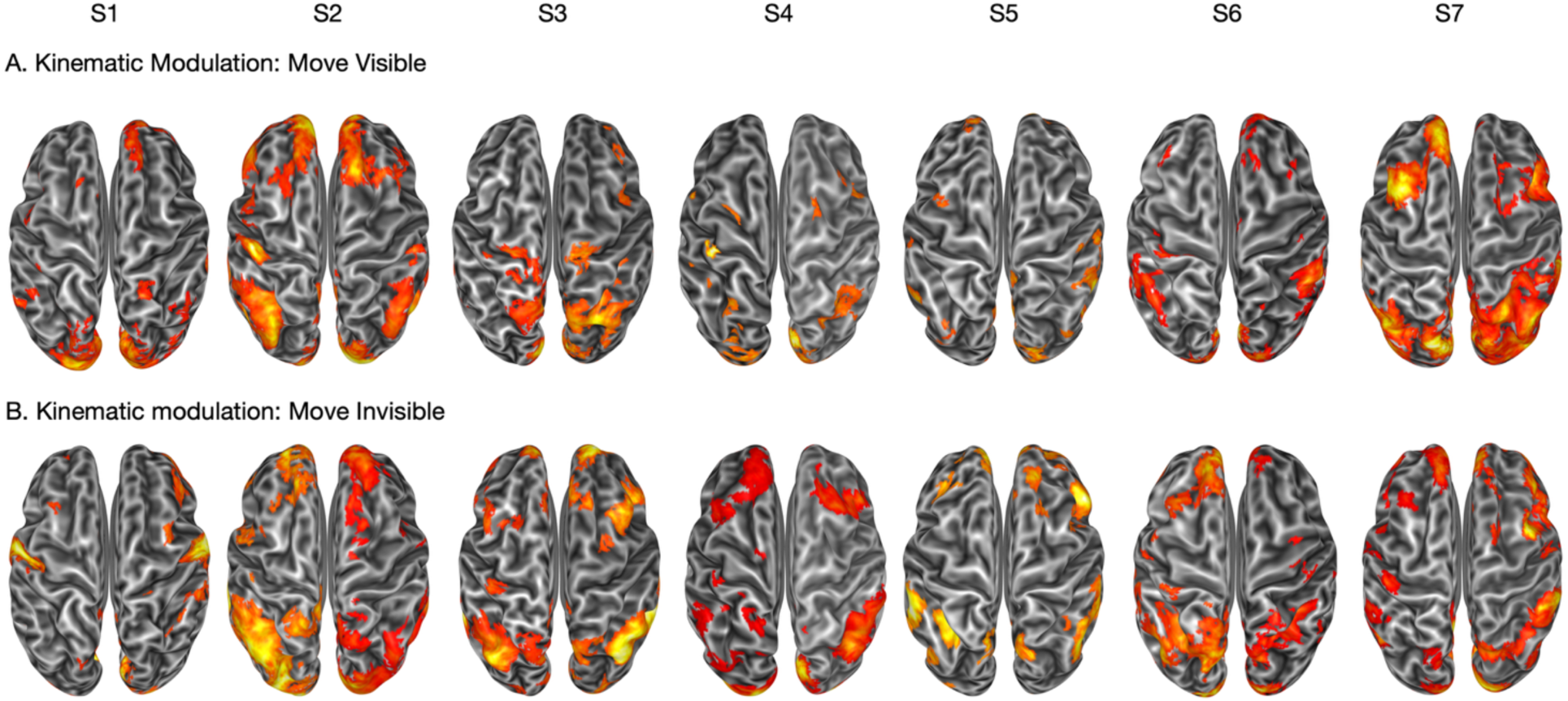
Effect of the kinematic parameters of the reaching movement on the individual brain activation of the 7 participants. For each participant, an inflated dorsal view of their surface reconstruction is shown, with regions showing a significant (p < 0.05 corrected for false discovery rate at the cluster level, with a cluster-forming threshold of p < 0.001 uncorrected) modulatory effect of the seven kinematic indices on the brain activation during movement trials with (A) or without (B) visual feedback. The seven kinematic indices were extracted from a group-level principal component analysis performed on a wide set of kinematic parameters (see text for details).

This differential pattern demonstrates that our approach successfully captures the impact of condition-specific kinematics on brain activity, highlighting the ability to differentiate parametric modulations across experimental conditions.

## 4. Discussion

This study introduces MOTUM, a novel system enabling real-time motion tracking during fMRI, integrated with immersive virtual reality. Our proof-of-concept experiment demonstrates that MOTUM effectively overcomes long-standing limitations in the neuroimaging of naturalistic motor behaviors, without compromising data quality or experimental control.

Traditional neuroimaging experiments have struggled to balance ecological validity with methodological rigor. Real-world actions are typically constrained or simulated using impoverished setups that strip away the complexity of natural movements. MOTUM addresses this gap by allowing participants to perform realistic, goal-directed actions— such as reach-to-grasp movements—within a virtual environment that mirrors real-life scenarios, all while simultaneously collecting high-resolution neural and kinematic data.

Despite the physical constraints of the MRI bore and potential issues with marker visibility, the system achieved robust motion tracking of arm, hand, and finger movements. Participants viewed and interacted with the virtual environment through a first-person avatar and reported strong feelings of agency and ownership, underscoring the quality of embodiment even in the absence of haptic feedback or physical object interaction.

A key concern in any movement-based fMRI paradigm is the potential for motion artifacts, mainly due to head displacement. Here, we found that reaching movements led to a modest increase in head movement, which remained well below critical thresholds, with only 0.1% of volumes exceeding the conventional task-fMRI threshold (FD > 0.5). This minimal data loss confirmed high signal quality and preserved adequate statistical power for our analyses, which also controlled for FD variance. We also evaluated the moment-by-moment hand and arm displacement effect on brain activity. While we did not find strong evidence of brain activity correlating with hand movements, arm displacement correlated with heightened activation of temporal, parietal, and frontal areas. We controlled for these sparse effects by regressing out hand and arm displacement in the following analyses.

A groundbreaking application of MOTUM is the possibility to disentangle the effect of kinematic movement properties on brain activity. Previous fMRI studies aiming at the same goal relied on contrasting conditions or performing repetition suppression analyses on the brain activity recorded during trials where a preset grip type or movement speed was required (e.g., Di Dio et al., 2013). Therefore, they did not investigate the natural, subtle trial-by-trial variation of the kinematic properties of the movements the participants performed. By providing continuous motion tracking of participants’ movements, MOTUM can pave the way to new possibilities. As a proof-of-concept in the present paper, we extracted latent dimensions of kinematic variation using principal component analysis (PCA). We showed that these modulate brain activity across sensorimotor and visuomotor networks. In particular, the AG, the pIPS, S1, MFG, area V6, and MT+ activity were modulated by kinematic movement properties.

While we did not focus on specific kinematics and their correlates in this analysis, our approach demonstrates that it is possible to identify specific impacts of individual kinematic indices on movement-related areas. For instance, previous studies have shown that movement velocity specifically affects motor, premotor, and superior parietal areas (Ferraina et al., 2009; Di Dio et al., 2013), while grip aperture selectively influences areas like the left aIPS, which scales with object size and grip type (Davare et al., 2007; Grol et al., 2007; Lehmann & Scherberger, 2015). Our methodology provides a framework that could capture such specific kinematic-neural relationships within a more comprehensive analysis of movement dynamics. Although preliminary, our results also point towards a different effect of kinematic movement properties on brain activity depending on the context, specifically, in our case, the absence/presence of visual feedback. While this has been previously hypothesized based on behavioral data (Jakobson & Goodale, 1991), the lack of human research with high-spatial resolution techniques such as fMRI makes this question worthy of further investigation.

Although our proof-of-concept focused on hand movements, MOTUM can be readily extended to other body parts, such as the lower limbs, which may offer even better tracking conditions given their visibility outside the scanner. This flexibility enables a broader range of real-life actions to be studied during fMRI, including locomotion and spatial navigation. Notably, the integration of virtual reality provides ecological realism and the ability to manipulate the environment and body representation in real time. This feature unlocks novel experimental designs—for instance, altering limb size or trajectory on the fly—to explore fundamental processes such as body ownership, visuomotor integration, and peripersonal space.

While the system is promising, some methodological limitations must be acknowledged. We restricted participation to individuals taller than 155 cm to minimize marker loss within the scanner bore and ensure sufficient arm visibility. Additionally, the reach-to-grasp task involved interaction with a virtual — rather than physical — object, as introducing a real object would have obstructed camera views. Although visual feedback was provided, our setup did not incorporate tactile or sensorimotor feedback. Future integration with amagnetic haptic devices could help overcome this limitation by enabling feedback upon virtual object contact.

In sum, we successfully implemented a unique real-time motion tracking system compatible with MRI. We demonstrated its effectiveness in capturing arm and hand movements without compromising data quality and in allowing a high-resolution mapping of motor dynamics onto neural data, offering the sensitivity to detect within-subject, between-trial effects. This system enables the investigation of naturalistic motor behaviors under controlled experimental conditions, offering a powerful new tool for motor neuroscience. MOTUM allows high-resolution, multi-joint kinematic recording in real time, enabling a detailed analysis of movement dynamics and their relationship with brain activity. This opens new avenues for studying motor control in ecologically valid settings. Moreover, the system supports standardized, reproducible data collection, which lays the groundwork for cross-laboratory comparisons and collaborative research on kinematic invariants (e.g., arm velocity) across tasks. In conclusion, MOTUM represents a significant methodological advancement, potentially transforming fMRI paradigms by bridging naturalistic motor behavior and brain imaging in a rigorously controlled yet flexible experimental framework.

## Data and Code Availability

All data and code will be available after acceptance of the paper based on request sent to the correspondence author with a possibility of needs for a formal data-sharing agreement, and approval from the requesting researcher’s local ethics committee.

## Author Contributions

Federica Bencivenga: Conceptualization, Formal Analysis, Methodology, Software, Visualization, Writing—Original Draft, Writing—Review and Editing.

Michelangelo Tani: Conceptualization, Data Curation, Formal Analysis, Investigation, Methodology, Software, Visualization, Writing—Original Draft, Writing—Review and Editing.

Krishnendu Vyas: Investigation.

Federico Giove: Data Curation, Funding Acquisition, Methodology, Resources. Steve Gazzitano: Investigation, Project administration, Resources.

Gaspare Galati: Conceptualization, Data Curation, Formal Analysis, Funding Acquisition, Methodology, Project administration, Software, Supervision, Writing—Review and Editing.

## Supplementary Material

**Supplementary Figure 1.**
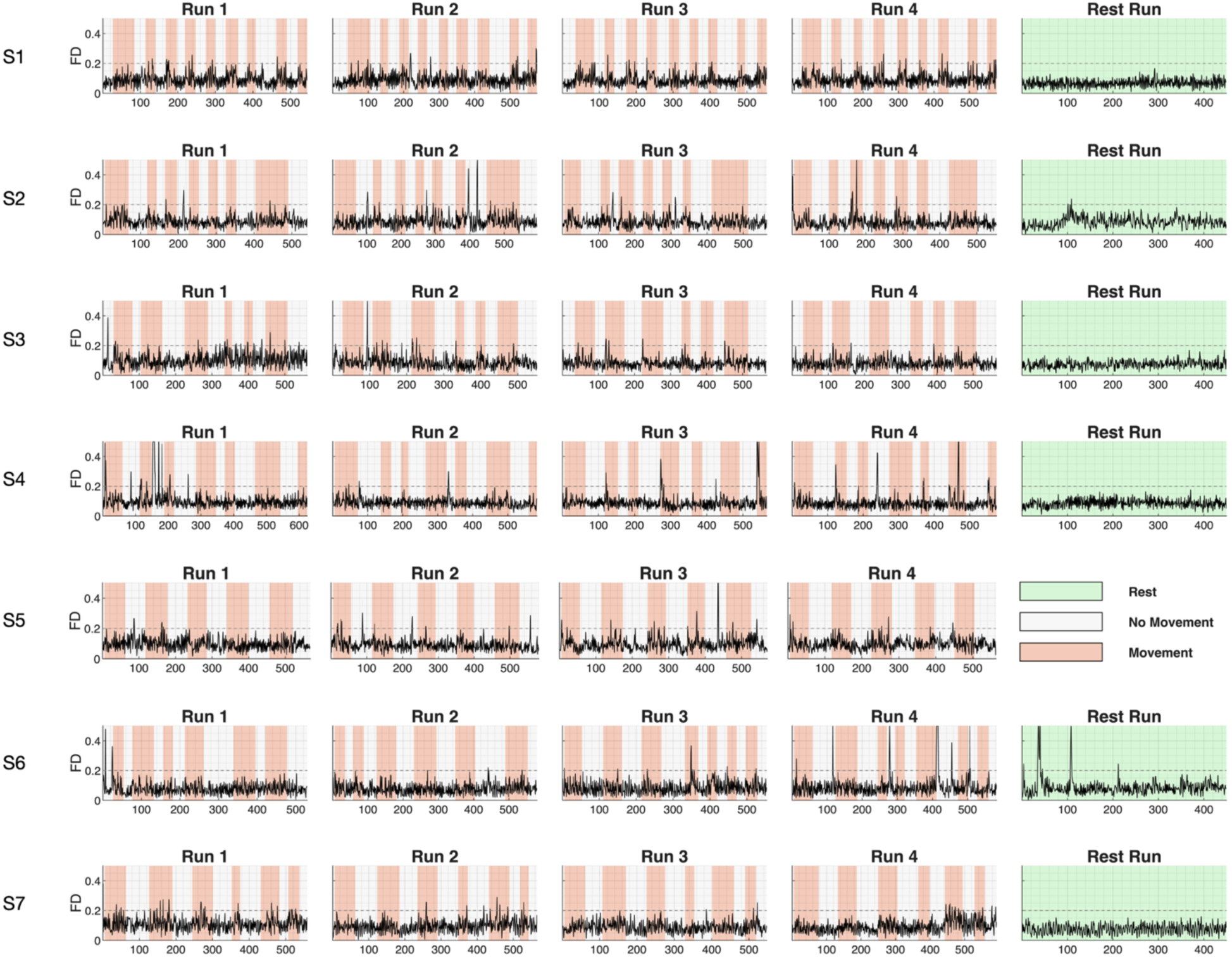
Framewise displacement (FD) timeseries across all scanning runs. Each row represents a different participant, with volume-wise FD values plotted on the y-axis (maximum 0.5 mm). Background colors indicate scan conditions: orange for movement blocks, white for no-movement blocks, and green for resting-state scans. The dashed horizontal line at 0.2 mm represents the optimal FD threshold reported in the literature, while 0.5 mm indicates the acceptable upper limit according to Siegel et al. (2014). Note that S5 did not complete the resting-state scan.

**Supplementary Table 1.**
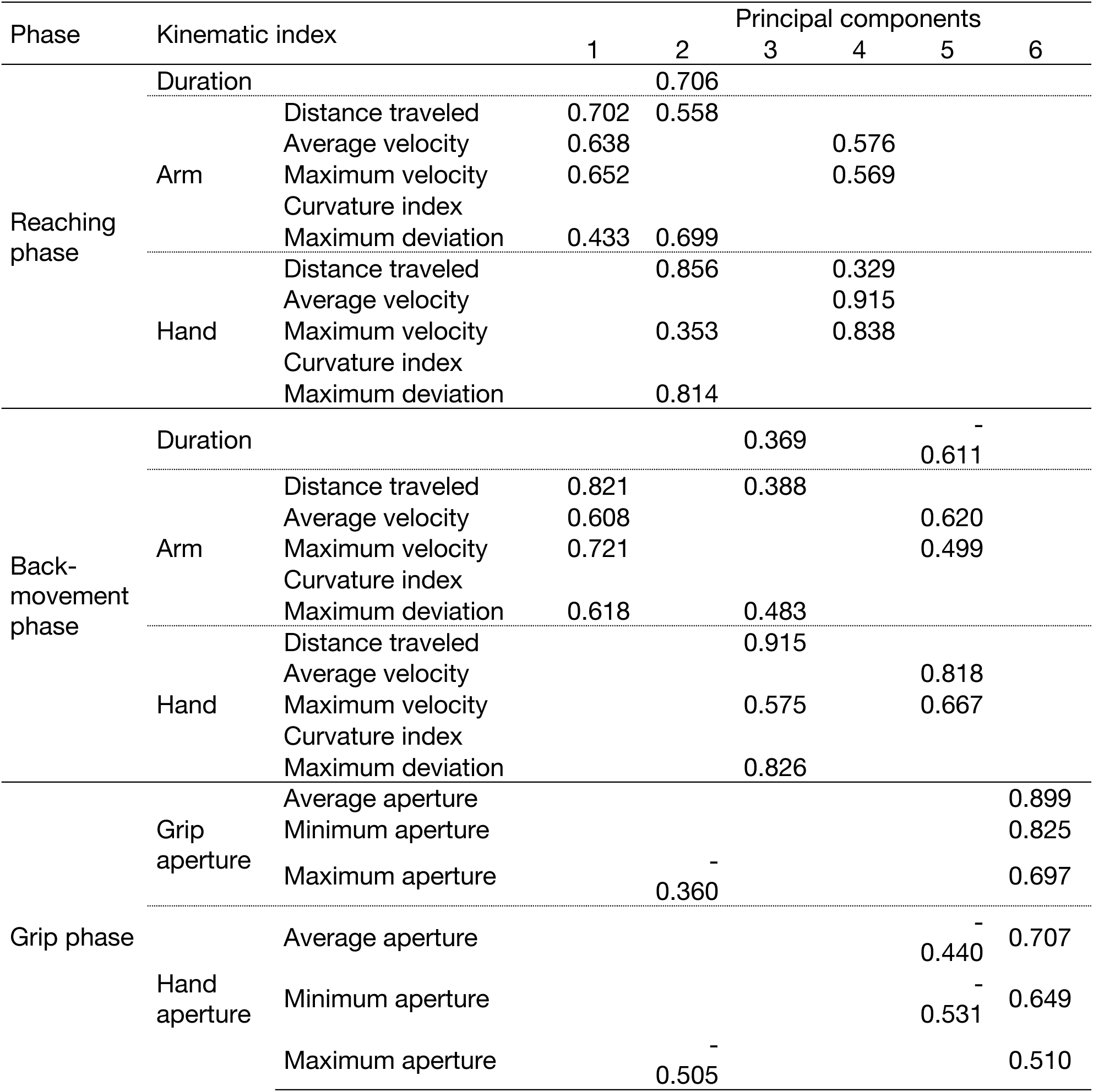
Loadings of the kinematic indices onto the extracted principal components.

